# Sensitivity analysis of voltage-gated ion channel models

**DOI:** 10.1101/2025.08.05.668838

**Authors:** Alon Korngreen

## Abstract

Markov models are widely used to describe the gating kinetics of voltage-gated ion channels, but increasing model complexity can introduce parameters that are difficult to constrain from macroscopic measurements. In this study, I used global variance-based Sobol sensitivity analysis to examine how model topology, stimulation protocol, and parameter uncertainty shape the accessibility of kinetic parameters in voltage-gated ion channel models. I analyzed progressively more complex Markov schemes, beginning with a two-state closed–open model, extending to a three-state linear closed–closed–open model, a cyclic three-state model with a direct closed–open transition, and a four-state closed–closed–open–inactivated model. Sensitivity was quantified for the open probability (Pₒ) under voltage-step and sinusoidal protocols. In linear models, parameter influence was concentrated in transitions directly connected to the open state, whereas distal closed–closed transitions contributed little to output variance. This weak influence was not rescued by higher-order interaction effects. Multi-frequency sinusoidal stimulation (20–200 Hz) preserved the same sensitivity hierarchy observed under step protocols, indicating that dynamic stimulation does not overcome the structural limitations of serial topologies. In contrast, introducing a cyclic pathway fundamentally redistributed sensitivity, showing that distal-parameter weakness is a consequence of serial arrangement rather than a universal property of Markov gating. Adding inactivation shifted dominant variance control to the open–inactivated transition during sustained depolarization, yet distal closed–closed transitions remained weak. Finally, constraining variability at the dominant activation edge shifted variance upstream, demonstrating that low Sobol indices reflect the relative flexibility of competing bottlenecks within a given ensemble rather than the intrinsic irrelevance of a transition. Together, these results define topology-dependent limits on parameter accessibility in Markov models of voltage-gated ion channels and provide practical guidance for selecting model complexity, designing informative protocols, and favoring cyclic over extended linear topologies when constructing robust channel models.

## Introduction

Voltage-gated ion channels are fundamental determinants of neuronal excitability. By shaping membrane conductances, they govern how neurons respond to synaptic input, generate action potentials, and regulate signal propagation across dendrites and axons (Koch, 1998; Koch and Segev, 2000; Roux et al., 2004). Because of this central role, ion channel models are a critical component of conductance-based descriptions of neuronal function and therefore contribute, indirectly but importantly, to the broader effort to understand how neurons process information and how neural circuits give rise to computation (Crick, 1989; Hausser et al., 2000; Crick and Koch, 2003; Spruston and Kath, 2004; Silver, 2010; Yuste and Church, 2014; Stuart and Spruston, 2015). At the same time, this broader connection should not obscure a more immediate and practical challenge: before ion channel models can support realistic neuronal simulations, their own internal structure and parameterization must be constrained in a way that is both biophysically meaningful and computationally tractable (Hay et al., 2011; Markram et al., 2015).

Neuronal models range from highly abstract representations that simplify neurons as point-like integrators to biologically detailed simulations that incorporate the full complexity of a neuron’s structure and ion channel biophysics (Koch and Segev, 2000; Poirazi and Mel, 2001; Poirazi et al., 2003). The latter, often called realistic models, have become increasingly feasible thanks to advances in experimental techniques and computational resources (Rapp et al., 1996; Roth and Häusser, 2001; Wood et al., 2004; Druckmann et al., 2007; Hines et al., 2008; Hendrickson et al., 2011; Schoen et al., 2012; Hay et al., 2013; Markram et al., 2015). Many of these models incorporate Hodgkin–Huxley-type formulations for voltage-gated channels, which use a compact set of gating variables to describe activation and inactivation kinetics (Hodgkin and Huxley, 1952). At the other end are Markov chain models, in which the channel is represented as a set of discrete conformational states connected by voltage-dependent transitions (Hille, 1992; Sakmann and Neher, 1995; Lampert and Korngreen, 2014; Sigg, 2014). Markov models offer substantially greater mechanistic flexibility and can represent features that are difficult to capture in Hodgkin–Huxley formulations, including multiple closed states, inactivated conformations, and alternative kinetic pathways. This flexibility, however, comes at a cost: as the number of states and transitions increases, so does the number of free parameters that must be inferred from experimental data (Almog and Korngreen, 2014, 2016). In practice, this often creates a mismatch between the model’s complexity and the information content of the measurements used to constrain it.

This problem is especially relevant when models are fit to macroscopic recordings. Macroscopic measurements provide rich information about the aggregate behavior of channel populations, but they do not directly report every microscopic transition in a Markov scheme (Nowotny et al., 2007; Fink and Noble, 2009). As a result, some parameters may have only a weak effect on the measured output, even though they remain part of the formal model structure. This creates a practical difficulty for optimization: parameters that exert little influence on the objective function are difficult to estimate reliably, can slow convergence, and may inflate model dimensionality without improving predictive value (Keren et al., 2005; Gurkiewicz and Korngreen, 2007; Menon et al., 2009; Hay et al., 2011; Ben-Shalom et al., 2012). The challenge is therefore not only to fit channel models, but also to determine which model topologies and which transition classes are actually accessible to a given type of measurement.

A related issue is that model topology itself may shape parameter accessibility. In a serial Markov chain, transitions that are topologically distant from the open state may contribute little to the variance of the measured output, whereas transitions directly coupled to the open state may dominate. If so, then the difficulty of estimating certain parameters is not merely numerical or algorithmic, but structural: it arises from the arrangement of states and transitions in the model. This possibility has important implications for how one should design Markov models. Adding states may increase formal realism, but it may also introduce parameters that are only weakly constrained by the available data. Conversely, alternative topologies, including cyclic arrangements, may redistribute sensitivity and improve the balance between model complexity and information content (Colquhoun et al., 2004; Menon et al., 2009).

In this study, I use global Sobol sensitivity analysis (Saltelli et al., 2008) to examine how parameter sensitivity depends on model topology, stimulation protocol, and parameter uncertainty in Markov models of voltage-gated ion channels. Rather than focusing on parameter fitting directly, I ask a more basic question: which transitions in a given model can substantially influence the variance of the open probability under commonly used voltage-clamp protocols? To address this, I analyze a progression of models of increasing complexity, beginning with a minimal two-state closed–open model, extending to linear three-state and four-state schemes that include inactivation, and then comparing these with a cyclic three-state topology. I examine both voltage-step and sinusoidal protocols and quantify not only first-order sensitivity but also total sensitivity and interaction contributions. I show that distal-parameter weakness in linear chains is a structural consequence of serial topology, that cyclic topologies fundamentally redistribute sensitivity, and that low Sobol indices reflect the relative flexibility of competing bottlenecks within a specified ensemble rather than the intrinsic irrelevance of a transition. Together, these results define topology-dependent limits on parameter accessibility and provide practical guidelines for selecting model complexity and topology when constructing robust and interpretable ion channel models.

## Methods

### Markov models of voltage-gated ion channels

In a standard Markov model for an ion channel with N states, the rate constants are represented by a conventional N×N rate matrix Q (Sakmann and Neher, 1995). In this matrix, each off-diagonal element represents the rate constant for the transition from state i to state j. Each diagonal element equals the negative sum of the off-diagonal elements in that row, ensuring that each row’s total is zero. If there is no direct transition from state i to state j, the corresponding element k_ij_ is set to zero. For voltage-gated channels, the simplest definition of the forward rate constant k_ij_ comprises a pre-exponential factor A_ij_. Additionally, an exponential factor, denoted z_ij_, is multiplied by the membrane potential, V. More complex functional representations for the voltage dependence of the rate constant have been applied (Destexhe, 2000; Destexhe and Huguenard, 2000). In this study, I used only this simple one.

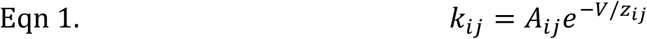

For forward rate constants, z_ij_ is negative, and for backward rate constants, z_ij_ is positive. This exponential factor can also depend on factors such as charge, pressure, temperature, and others (Hille, 1992). At any given moment, a state probability vector P, which has a dimension of N, represents the occupancy of the N states in the model. The Kolmogorov differential equation describes the dynamics of a Markov process (Sigg, 2014).

The time evolution of state occupancies in the Markov models was described using the Kolmogorov equation. At any given moment, the occupancy of the N states is represented by a state probability vector **P**(t) = [P₁(t), P₂(t),…, Pₙ(t)], where Pᵢ(t) denotes the probability that the channel occupies state *i* at time *t*. Transitions between states are governed by the voltage-dependent transition rate matrix Q(V), defined above, in which each off-diagonal element Qᵢⱼ(V) corresponds to the transition rate from state *i* to state *j*, and diagonal elements are defined such that each row sums to zero ensuring conservation of total probability. The temporal evolution of state occupancies is therefore given by

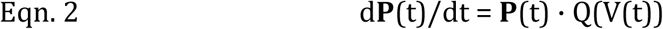

where V(t) is the imposed voltage-clamp command. This formulation assumes that channel gating follows a continuous-time Markov process in which transitions depend only on the current state and the instantaneous membrane potential.

Transition rates were voltage-dependent and followed the exponential formulation described above, with each element of Q(V) defined by the corresponding voltage-dependent rate constant kᵢⱼ(V). Numerical integration of the Kolmogorov equation yielded the time-dependent state occupancies for each simulation. The model output used for the sensitivity analysis was the open probability, computed as the sum over all states corresponding to the open conformation (i.e., the open-state occupancy for single-open-state models). Simulations were initialized at steady-state occupancy at the holding potential.

The Markov state-probability dynamics were integrated by solving Eqn. 2 with SciPy’s solve_ivp, using an implicit stiff solver (BDF by default, with Radau as an alternative). Default solver tolerances of rtol = atol = 10⁻⁶ were applied throughout. Initial conditions were defined as the stationary distribution at t=0, obtained from the eigenvector of Q^T^ associated with the eigenvalue closest to zero and normalized to unit mass. To monitor numerical quality, I recorded the maximum mass-conservation error 𝐦𝐚𝐱𝑡 ∣ ∑_𝒊_ 𝒙_𝒊_ (𝑡) − 1 ∣, probability-bound violations (states outside [0,1]), generator row-sum residuals, microscopic reversibility residuals (when applicable), and an optional stiffness-consistency metric, defined as the maximum absolute difference between BDF and Radau solutions on the same output grid.

### Sensitivity analysis

Sensitivity analysis evaluates how uncertain parameters influence the variability of model outputs. Numerous sensitivity measures are available (Hamby, 1994; Saltelli et al., 2008; Zi, 2011). In this study, I employed a global variance-based sensitivity analysis that calculates Sobol sensitivity indices, a well-established approach (Sobol, 2001). This global and non-intrusive method allows us to investigate interactions among model parameters (Zi, 2011). Interactions occur when the combined effects of parameters on the output are non-additive.

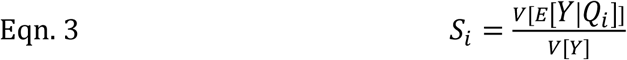

Several Sobol’ indices exist, with the first-order Sobol sensitivity index (*S_i_*) being the most commonly used. S_i_ measures the direct influence of each parameter on the model output variance. 𝐸[𝑌|𝑄_𝑖_] represents the expected value of the output Y when the parameter Q_i_ is held constant. 𝑉[𝐸[𝑌|𝑄_𝑖_]] quantifies the variance explained by Q_i_ alone, and V[Y] is the variance of the output. The first-order Sobol sensitivity index indicates how much the model’s variance is expected to decrease when parameter Q_i_ is held constant. For example, if S_i_ =0.3, then 30% of the output variance is explained by Q_i_ alone. The sum of all first-order Sobol sensitivity indices cannot exceed one and will only equal one if there are no interactions between parameters (Glen and Isaacs, 2012).

In this study, I also calculated the total sensitivity indices.

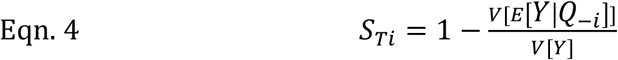

𝐸[𝑌|𝑄_−𝑖_] represents the expected value of the output Y when all parameters, except Q_i,_ are held constant. 𝑉[𝐸[𝑌|𝑄_−𝑖_]] quantifies the variance explained by all variables except Q_i_. For example, if S_Ti_ =0.3, then Q_i_ (including its interactions with other variables) explains 30% of the output variance.

All simulations were carried out using custom Python code and the SALib sensitivity analysis library (Herman and Usher, 2017). Sobol indices were estimated using the Saltelli sampling scheme with a base sample size of N = 1024, yielding 6,144 model evaluations for four-parameter models and 10,240 for eight-parameter models; this sample size was chosen to provide stable index estimates across all topologies and protocols while keeping computational cost tractable. Convergence was confirmed by repeating key analyses at N = 512 and N = 2048.

The code is publicly available on GitHub at https://github.com/alon67/MarkovSensitivity.

## Results

The simplest Markov model for an ion channel consists of two states: closed and open (Scheme 1). For a voltage-gated channel, the rate constants describing the transitions between these states are voltage-dependent, as detailed in the Methods section. This straightforward formulation contains four free parameters: two pre-exponential rate constants (A₁₂ and A₂₁) and two voltage-dependence parameters (z₁₂ and z₂₁). To evaluate how variations in these parameters influence model behavior, I examined the sensitivity of the open probability (P_o_) under a voltage step from −80 mV to +10 mV.

**Scheme 1.**
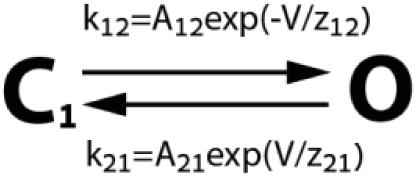

In this study, sensitivity analysis is performed on the open probability rather than on macroscopic current. For a voltage-step protocol, macroscopic current can be written as 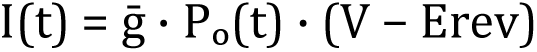

Within a fixed-voltage epoch, both the maximal conductance 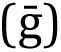 and the driving-force term (V − E_rev_) are constants, so current is a constant multiple of Pₒ. Consequently, for step protocols, sensitivity rankings for current within these epochs are expected to match those obtained for Pₒ, up to a uniform rescaling of the output variance. Under time-varying voltage commands, the multiplicative driving force can reshape the current’s sensitivity. Thus, I limited the scope of this manuscript to investigating the impact of parameter variability on P_o_.

Figure 1A shows representative simulations of P_o_ (thin black traces) together with the population average (thick red trace). Parameter values were sampled across prescribed ranges, producing a broad diversity of activation and deactivation kinetics. Despite this variability, the ensemble mean displayed a clear activation plateau during depolarization and a rapid return toward baseline upon repolarization. This plateau region provided a natural reference point for interpreting the model’s sensitivity structure.

**Figure 1.**
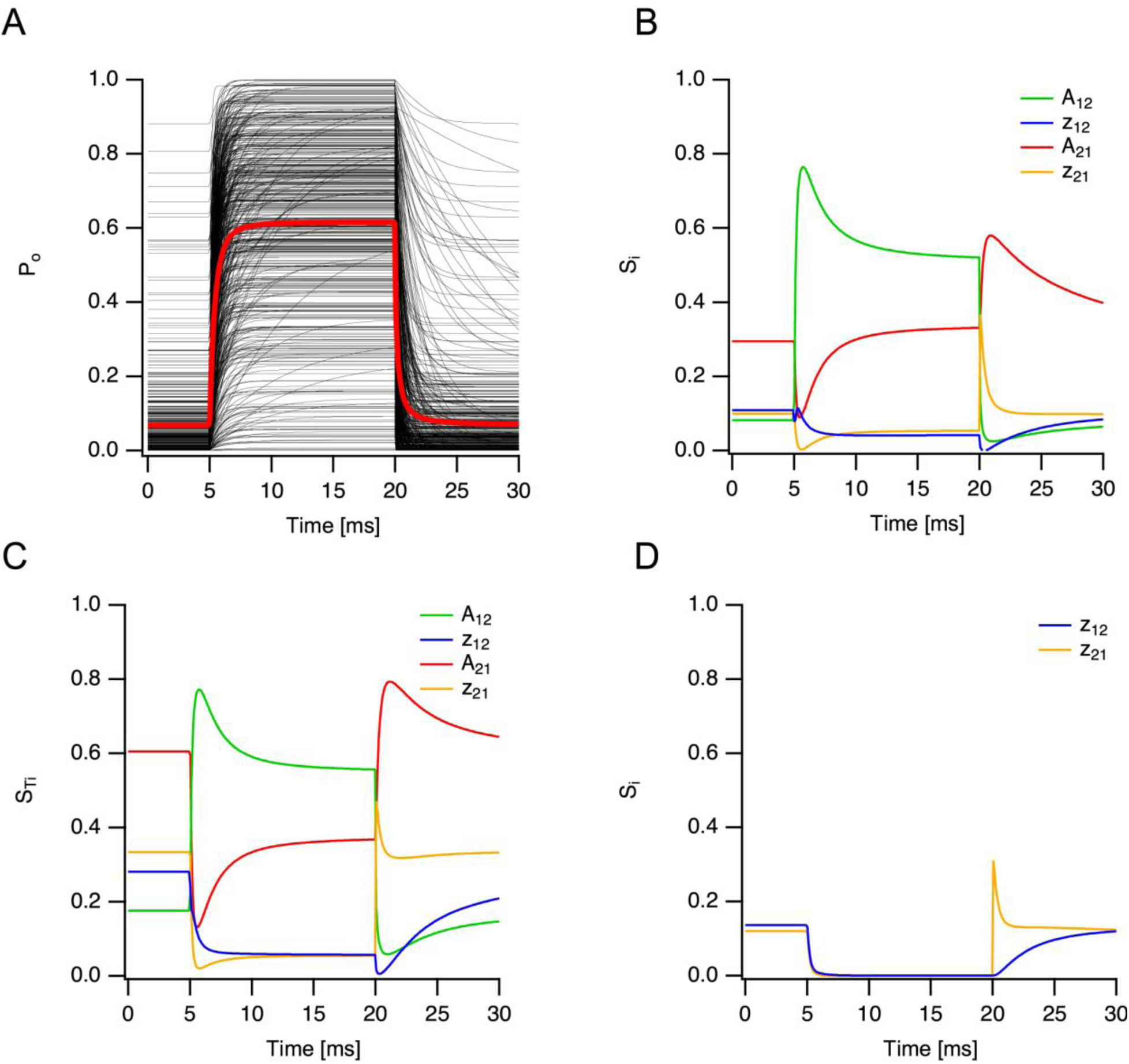
Sensitivity analysis of a two-state Markov model of a voltage-gated ion channel under a step protocol. (A) Representative simulations (thin black traces) and population average (thick red trace) of the open probability (P_o_) in response to a voltage step from −80 mV to +10 mV. Parameters (A₁₂, A₂₁: 0.0001–1 ms⁻¹; z₁₂, z₂₁: 1–80 mV) were sampled within the specified ranges. Simulations were initialized at a holding potential of −80 mV. (B) Time-resolved first-order Sobol indices (Sᵢ) for the four free parameters: forward rate amplitude (A₁₂, green), forward voltage dependence (z₁₂, blue), backward rate amplitude (A₂₁, red), and backward voltage dependence (z₂₁, yellow). (C) Time-resolved total Sobol indices (S_Ti_) for the same parameters and protocol. (D) Sanity check. First-order indices for z₁₂ (blue) and z₂₁ (yellow) during a step from −80 mV to 0 mV. As expected, sensitivity to the voltage-dependence parameters vanishes at 0 mV. Sobol analysis used Saltelli sampling with N = 1024, yielding 6144 model evaluations per time point.

The time-resolved first-order Sobol indices (Sᵢ) are shown in Figure 1B. At the holding potential (−80 mV), the model exhibited modest sensitivity primarily to the backward rate constant (A₂₁), reflecting the dominance of the closed state at rest. Upon depolarization, sensitivity shifted sharply toward the forward transition (A₁₂), consistent with the increased probability of channel opening. The voltage-dependence parameters (z₁₂ and z₂₁) displayed transient sensitivity peaks during the activation and deactivation phases, corresponding to periods when the exponential voltage terms most strongly influenced the rate constants.

Total Sobol indices are shown in Figure 1C. As expected for a four-parameter system, values were slightly larger than the corresponding first-order indices, indicating the presence of modest interaction effects. However, the overall ranking of parameter influence remained unchanged, with transitions directly governing opening and closing dominating the variance of P_o_.

To validate the voltage-dependent formulation and the implementation of the sensitivity analysis, I performed a control simulation in which the membrane potential was stepped from −80 mV to 0 mV (Figure 1D). At 0 mV, the exponential voltage-dependent component of the rate constants is nullified. As predicted, the sensitivity indices for the voltage-dependence parameters (z₁₂ and z₂₁) collapsed to zero under this condition, confirming that the analysis correctly captures the functional structure of the rate equations.

To assess the sensitivity analysis’s dependence on the assumed sampling distribution, I repeated the simulation of the two-state model using both uniform and log-uniform parameter sampling. As expected for a variance-based method, the absolute magnitudes of the Sobol indices changed under the two sampling schemes. In particular, log-uniform sampling increased the dominance of the forward prefactor term and reduced the contribution of the backward prefactor term relative to uniform sampling. However, the qualitative sensitivity hierarchy was unchanged: in both cases, the rate prefactors remained the dominant contributors, whereas the voltage-sensitivity terms remained weak. Thus, for the two-state model, the choice of sampling distribution affected the quantitative partitioning of variance but did not alter the mechanistic interpretation of the results. These data are not shown. For completeness, the publicly available code includes an option to switch between uniform and log-uniform sampling, allowing the same comparison to be examined for all models analyzed in this study.

Together, these results establish the baseline sensitivity structure of the simplest Markov representation. Under standard step stimulation, parameter influence follows the expected activation–deactivation asymmetry, and interaction effects are limited. This minimal model provides a controlled reference point for examining how increasing topological complexity alters the relationship between parameter dimensionality and observable sensitivity.

To move beyond the minimal two-state formulation, I examined parameter sensitivity in a three-state linear Markov model composed of two closed states and one open state (Scheme 2). The model was stimulated using the same step protocol described above. Representative simulations of P_o_ are shown in Figure 2A. The additional closed state broadened the diversity of responses across parameter sets, yet the population average exhibited a well-defined activation plateau during depolarization. Sensitivity indices were therefore quantified over the steady-state activation window (16–19 ms), corresponding to the plateau phase of the depolarizing step.

**Figure 2.**
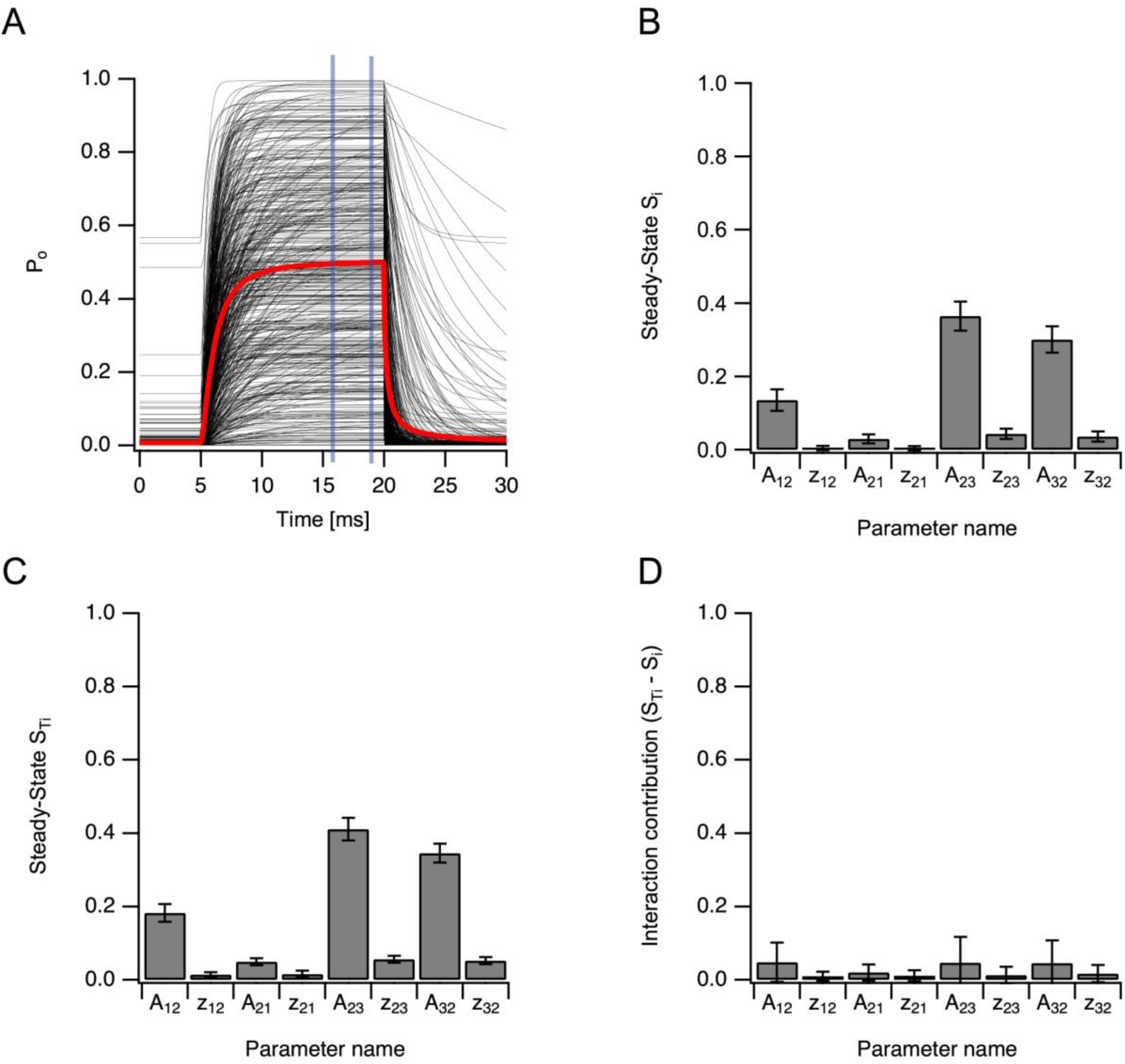
Quantitative steady-state sensitivity structure of a three-state linear Markov model. (A) Representative simulations (thin black traces) and population average (thick red trace) of the open probability (P_o_) in response to a voltage step from −80 mV to +10 mV. Simulations were initialized at steady state at −80 mV. The steady-state activation window used for quantitative analysis (16–19 ms) lies within the plateau phase of depolarization (indicated by vertical lines). (B) Steady-state first-order Sobol indices (Sᵢ) for all free parameters of the three-state linear model, computed as the mean Sᵢ over the activation plateau (16–19 ms). Error bars indicate bootstrap confidence intervals. Forward transitions directly coupled to the open state exhibit the largest contributions to output variance, whereas parameters governing closed–closed transitions contribute minimally. (C) Steady-state total Sobol indices (S_Ti_) over the same window. The total index includes both direct effects and interaction contributions. (D) Interaction contribution, defined as S_Ti_ − Sᵢ, quantifying the variance attributable to parameter interactions. Interaction effects are modest and do not substantially alter the ranking of parameter influence, indicating that low first-order sensitivity reflects a weak contribution rather than a hidden interaction structure. Sobol analysis used Saltelli sampling with N = 1024, yielding 10,240 model evaluations per time point. Confidence intervals were estimated by bootstrap resampling.

**Scheme 2.**
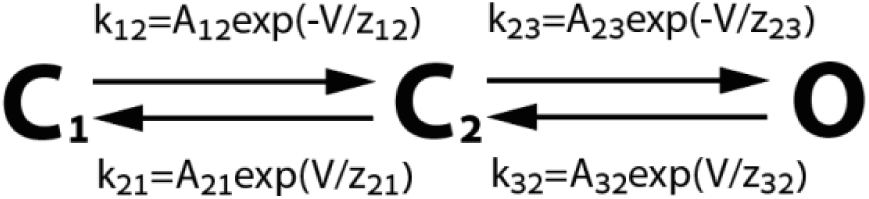

A clear hierarchy of parameter influence emerged (Figure 2B). Transitions directly coupled to the open state dominated the variance of P_o_, with the forward transition into the open state (A₂₃) exhibiting the largest contribution and the backward transition from the open state (A₃₂) contributing substantially. In contrast, parameters governing transitions between closed states displayed consistently low first-order sensitivity. Thus, even in a slightly more complex topology, the steady-state behavior of the model remains overwhelmingly controlled by rates directly linked to the open conformation.

Although S_Ti_ values were uniformly higher than Sᵢ (Figure 2C), reflecting the inclusion of interaction effects, the ranking of parameter importance was preserved. To evaluate whether low first-order sensitivity could be compensated by higher-order interactions, I explicitly computed the interaction contribution (S_Ti_ − Sᵢ, Figure 2D). Interaction effects were modest across all parameters and remained small for closed–closed transitions. Parameters that were weak at the first-order level did not acquire substantial influence through interactions.

These results demonstrate that the diminished influence of distal closed-state transitions in linear Markov models is not an artifact of neglected interaction terms but a structural property of the topology under standard step stimulation. Adding additional closed states increases parameter dimensionality without proportionally increasing the capacity of macroscopic P_o_ measurements to constrain those parameters. Consequently, extended linear chains introduce degrees of freedom that remain weakly informed by steady-state activation data, reinforcing the need to reconsider topology when designing Markov models of voltage-gated ion channels.

To determine whether dynamic stimulation alters the sensitivity structure observed under step protocols, I next examined the three-state linear model under sinusoidal voltage driving. Sinusoidal voltage protocols have been proposed as an efficient strategy for rapid characterization of ion channel kinetics and parameter fitting (Beattie et al., 2018), motivating the present test of whether oscillatory stimulation alters the sensitivity hierarchy of the linear model. The membrane potential was varied sinusoidally (±60 mV, centered at 0 mV). Representative simulations of open probability (Pₒ) at 50 Hz are shown in Figure 3A. The model exhibits stable oscillatory gating behavior across the sampled parameter space, with the population average tracking the sinusoidal command without evidence of refractory dynamics. These simulations confirm that the model operates in a fully dynamic regime, distinct from the step protocol previously used.

**Figure 3.**
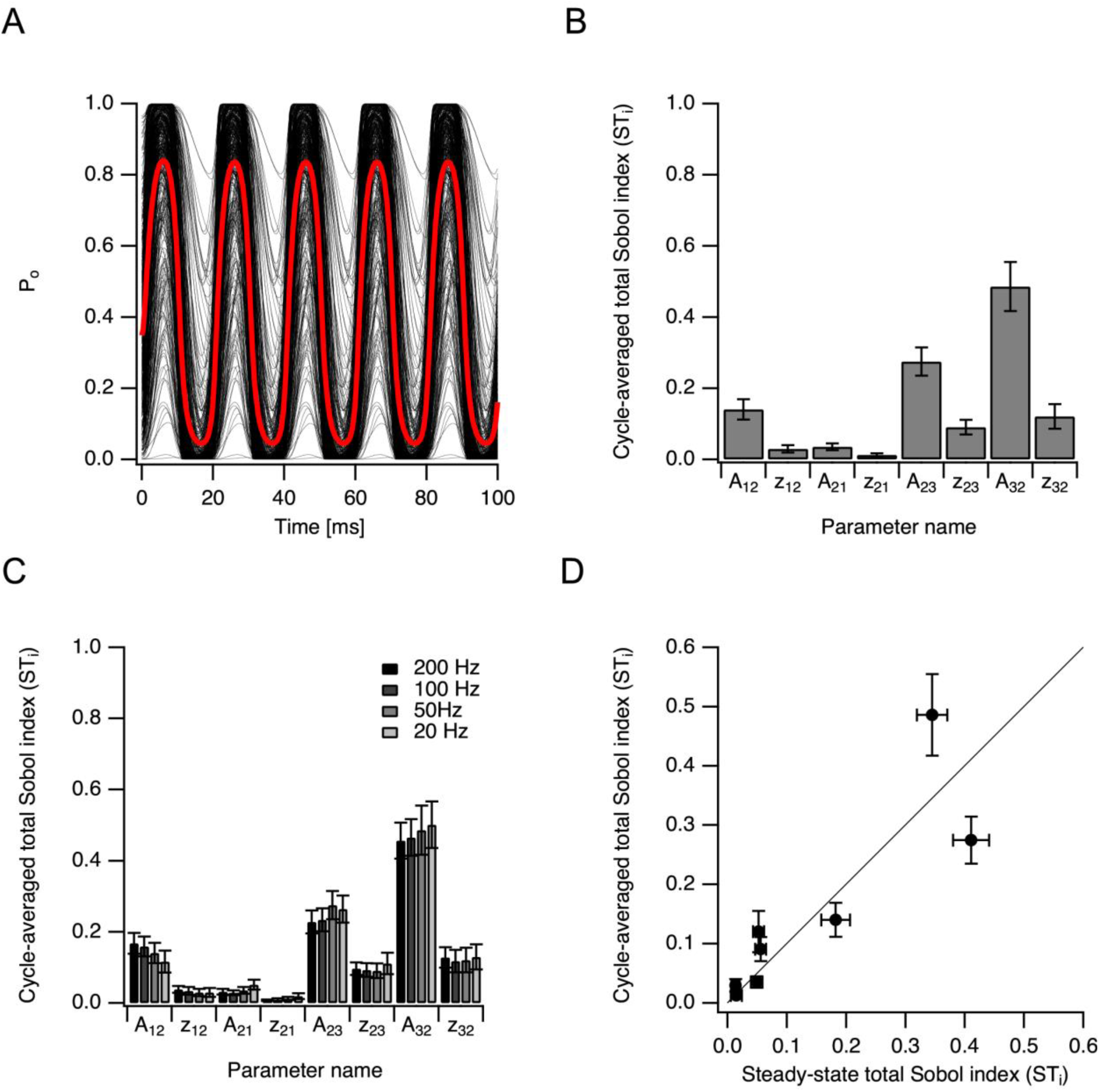
Dynamic stimulation does not alter the structural hierarchy of parameter sensitivity. (A) Representative simulations of open probability (Pₒ) under sinusoidal voltage stimulation (±60 mV, centered at 0 mV, 50 Hz). Thin gray traces show individual simulations across the sampled parameter space (Saltelli sampling, N = 1024); the thick red trace shows the population average. The model exhibits stable oscillatory gating dynamics without refractory behavior. (B) Cycle-averaged total Sobol indices (S_T_ᵢ) at 50 Hz. Sensitivity indices were computed at each time point and averaged over the full sine cycle. Points denote mean STᵢ values; error bars indicate bootstrap confidence intervals. Transitions directly connected to the open state dominate output variance, whereas distal transitions remain weak. (C) Frequency dependence of cycle-averaged S_T_ᵢ across 20–200 Hz (±60 mV, offset 0 mV). Parameter ranking remains stable across stimulation frequencies, indicating that oscillatory dynamics do not redistribute sensitivity toward weak transitions. (D) Comparison of steady-state (step protocol, 16–19 ms window) and sine (50 Hz) cycle-averaged total Sobol indices. Each point represents one parameter; horizontal and vertical error bars indicate bootstrap confidence intervals for the step and sine conditions, respectively. The solid line denotes the identity line. Parameter ranking is strongly preserved between protocols (Spearman ρ = 0.93, p = 8.6 × 10⁻⁴), demonstrating that dynamic stimulation does not rescue structurally weak parameters in the linear topology. All analyses were performed with base sample size N = 1024 (D = 8 parameters; 10,240 model evaluations per protocol). Only total Sobol indices are shown.

To quantify parameter influence under oscillatory driving, S_T_ᵢ were computed at each time point and averaged over the full stimulation cycle. Figure 3B shows the cycle-averaged STᵢ values at 50 Hz. The sensitivity hierarchy closely mirrors that observed under steady-state activation: transitions directly connected to the open state dominate output variance, while distal transitions remain weak. Bootstrap confidence intervals were small relative to the magnitude of dominant indices, indicating stable estimation. Thus, oscillatory stimulation does not elevate the influence of parameters that were previously weak under step conditions.

To assess robustness to stimulation frequency, the analysis was also repeated at 20, 100, and 200 Hz (Figure 3C). For each frequency, S_T_ᵢ values were computed at every time point and averaged over the full sine cycle. Across this fourfold range of frequencies, the ordering of parameter importance remained remarkably stable. Although individual S_T_ᵢ magnitudes shifted modestly with frequency, the same subset of transitions consistently accounted for the majority of output variance. No frequency selectively enhanced the contribution of distal closed-state transitions.

Finally, to quantify the similarity between the dynamic and steady-state sensitivity structures, cycle-averaged S_T_ᵢ values at 50 Hz were directly compared with steady-state STᵢ values obtained from the step protocol. Each parameter is represented as a single point in Figure 3D, with horizontal and vertical error bars reflecting bootstrap confidence intervals for the step and sine conditions, respectively. The solid line denotes the identity line. Parameter ranking was strongly preserved between protocols (Spearman ρ = 0.93, p = 8.6 × 10⁻⁴). Even when accounting for uncertainty through inverse-variance weighting, rank correlation remained high (ρ_w_ = 0.84).

Together, these results demonstrate that oscillatory voltage stimulation does not fundamentally redistribute parameter influence in the linear topology. The sensitivity hierarchy observed under steady-state activation persists across dynamic driving and across stimulation frequencies. Thus, weak parameters remain weak not because of a specific stimulation protocol, but because of the structural organization of the linear Markov chain.

To determine whether the sensitivity hierarchy observed in the linear chain reflects a structural constraint of the topology rather than a generic feature of three-state models, I next examined a cyclic three-state mechanism (Scheme 3). In this topology, a direct transition between C₁ and O was introduced, closing the loop. Microscopic reversibility was enforced by constraining the rate constants along the cycle, reducing the number of independent parameters relative to the unconstrained formulation (Colquhoun et al., 2004). This removed a pair of parameters (A_32_ and z_32_) from the sensitivity analysis since this transition rate constant was calculated from the thermodynamic cycle constraint of all other transitions. All simulations were performed under identical conditions to the linear model (+10 mV step; steady-state window 16–19 ms; Saltelli sampling, N = 1024).

**Scheme 3.**
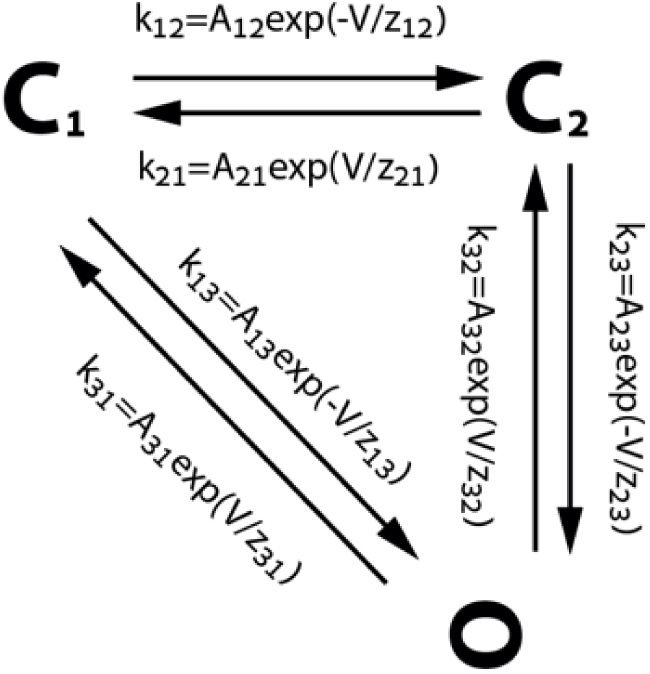

In contrast to the linear CCO model (Figure 2), where sensitivity was concentrated on the C₂–O transition, the cyclic topology exhibits a different hierarchy (Figure 4A). Sensitivity is now dominated by the direct C₁–O transitions, while the previously dominant C₂–O transition (A₂₃, z₂₃) collapses to near-zero influence. The voltage sensitivities associated with the C₂–O transition are similarly reduced. Thus, the addition of a direct C₁–O transition alters the location of variance control within the network.

**Figure 4.**
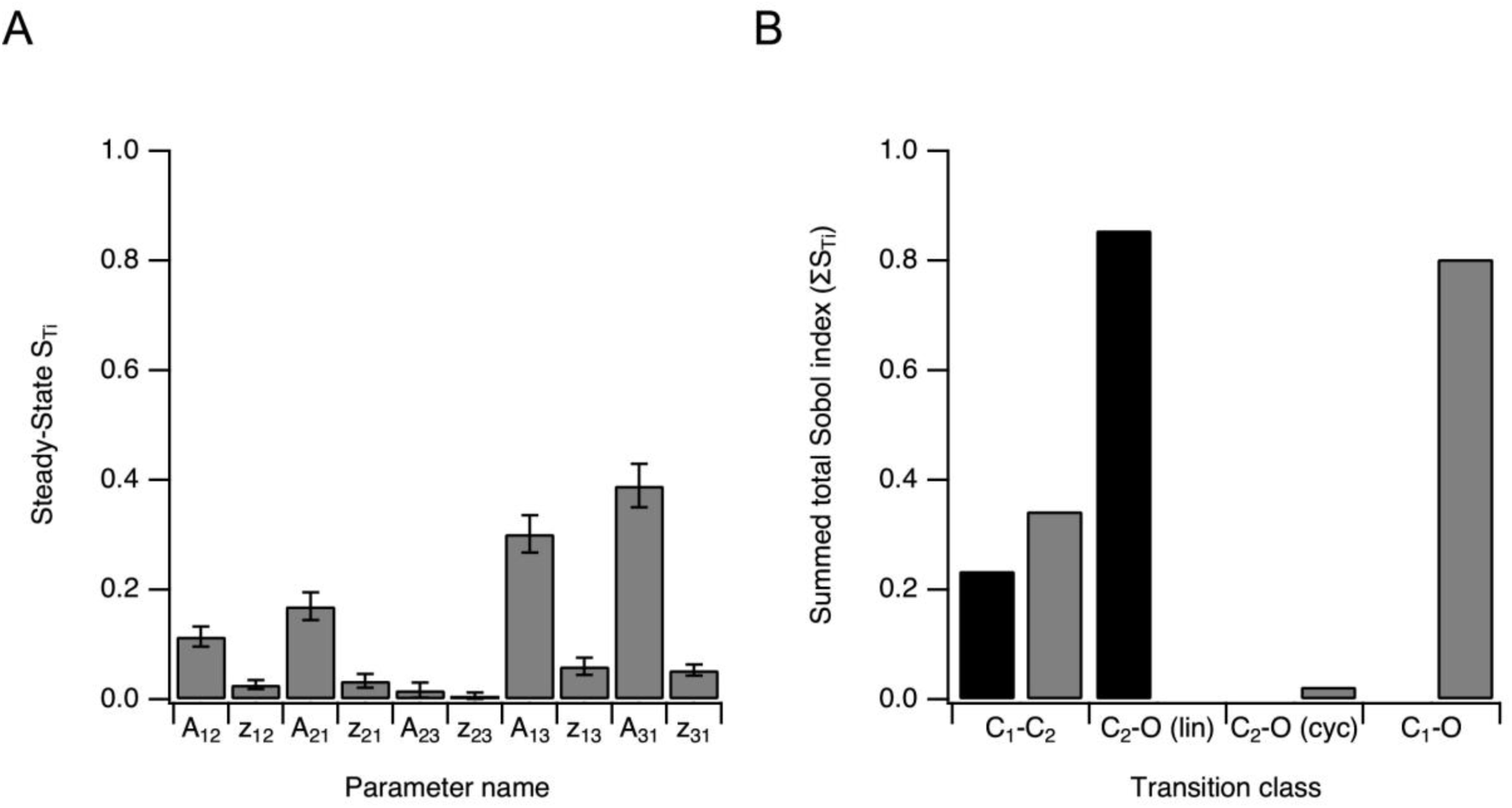
Cyclic topology redistributes steady-state parameter sensitivity. (A) Steady-state total Sobol indices (S_T_ᵢ) for the cyclic mechanism (Scheme 3) at +10 mV (analysis window 16–19 ms). Bars denote mean S_T_ᵢ values; error bars indicate bootstrap confidence intervals (Saltelli sampling, N = 1024). Parameters are shown in structural order corresponding to the transition classes C₁–C₂ (A₁₂, z₁₂; A₂₁, z₂₁), C₂–O (A₂₃, z₂₃), and C₁–O (A₁₃, z₁₃; A₃₁, z₃₁). In contrast to the linear chain (Figure 2), sensitivity is no longer concentrated on the C₂–O bottleneck. (B) Redistribution of sensitivity across transition classes. Bars show the summed total Sobol index (ΣSTᵢ) within each transition class. Black bars correspond to the linear CCO model; gray bars correspond to the cyclic model (Scheme 3). Transition classes are indicated on the x-axis as C₁–C₂, C₂–O (lin), C₂–O (cyc), and C₁–O. In the linear model, sensitivity is dominated by the C₂–O bottleneck. In the cyclic topology, variance shifts to the direct C₁–O pathway, while the contribution of the middle C₂–O edge is markedly reduced. Thus, closing the loop structurally relocates sensitivity rather than merely redistributing it within the original chain. All simulations were performed under identical sampling conditions as in Figures 2 and 3.

To quantify this redistribution at the transition level, I aggregated S_T_ᵢ values within each edge class (Figure 4B). In the linear model, the summed sensitivity (ΣS_T_ᵢ) is overwhelmingly concentrated in the C₂–O transition, consistent with a bottleneck structure. In the cyclic model, this pattern is reversed; ΣS_T_ᵢ shifts to the C₁–O transition, while the contribution of the middle C₂–O transition is markedly diminished. The C₁–C₂ transition exhibits a modest increase relative to the linear case but does not dominate the variance. Importantly, this redistribution occurs despite identical voltage protocols and sampling strategies, indicating that the shift arises from topology rather than stimulation conditions.

These results demonstrate that the “distal parameter weakness” observed in the linear chain is not an intrinsic property of three-state gating models, nor a consequence of the steady-state protocol. Rather, it reflects the structural constraint imposed by a serial topology in which variance must flow through a single open-adjacent bottleneck. Closing the loop with a direct C₁–O pathway removes this constraint and relocates sensitivity accordingly. The cyclic mechanism, therefore, reveals that parameter influence in Markov models is determined primarily by topological organization, with microscopic reversibility ensuring thermodynamic consistency but not preventing redistribution of control.

Repeating the cyclic-model analysis under log-uniform sampling altered the absolute Sobol magnitudes and modestly changed the balance among the dominant prefactor terms, but it did not change the qualitative conclusion. In both cases, the direct C₁–O pathway remained the dominant source of variance, whereas the intermediate C₂–O edge remained weak. Thus, the topological redistribution induced by closing the loop is robust to the choice of sampling distribution (data not shown).

Following the analysis of cyclic topology (Figure 4), I next asked whether introducing inactivation alters the structural sensitivity hierarchy observed in linear chains. To address this, I extended the model to include an inactivated state in a linear C₁–C₂–O–I configuration (Scheme 4). The system was stimulated with a prolonged depolarizing step to +10 mV lasting 80 ms within a 100 ms simulation, allowing activation and inactivation dynamics to unfold. Sensitivity indices were computed using Saltelli sampling (N = 1024), and steady-state values were averaged over the 65–75 ms window during sustained depolarization.

**Scheme 4.**
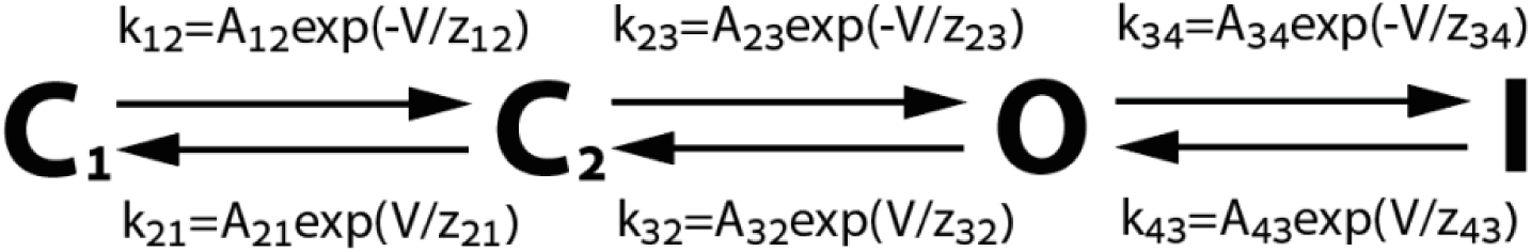

Representative Pₒ traces (Figure 5A) display rapid activation followed by a gradual decline reflecting entry into the inactivated state. The ensemble average confirms that the model captures physiologically plausible activation–inactivation behavior under sustained depolarization.

**Figure 5.**
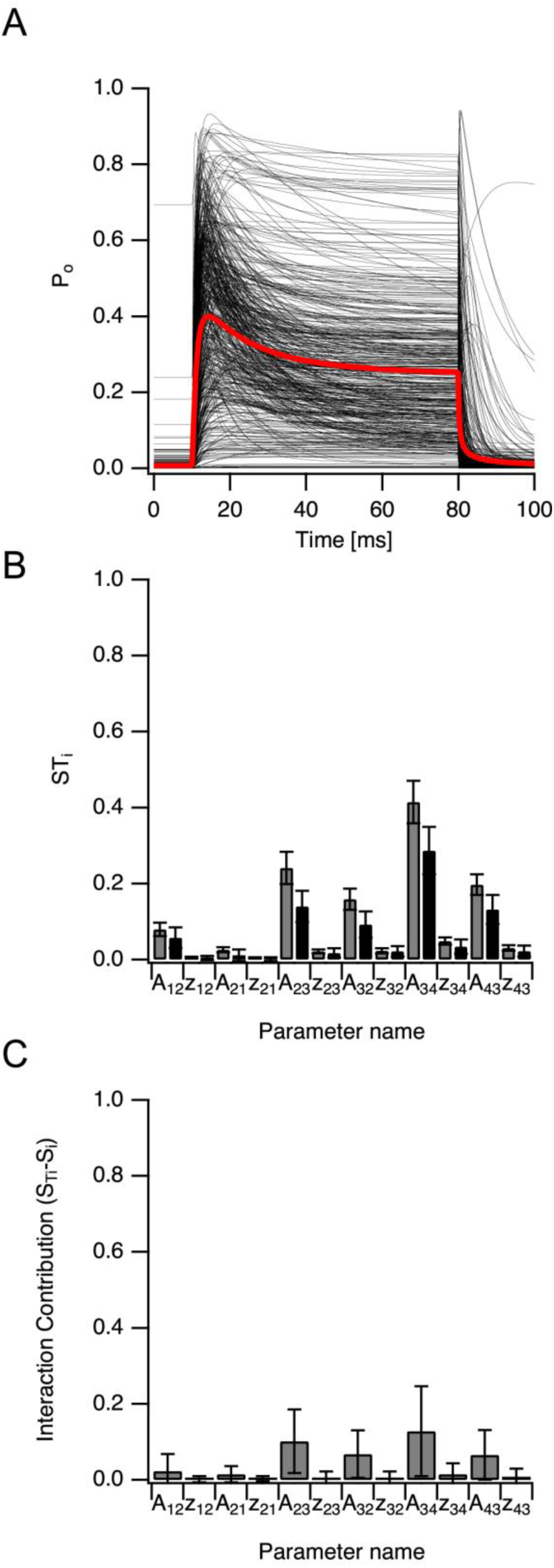
Inclusion of inactivation reshapes sensitivity and reveals interaction structure under sustained depolarization. (A) Open probability (Pₒ) traces for the linear C₁–C₂–O–I model (Scheme 4) under a prolonged depolarizing step. The membrane potential was held at baseline for 10 ms, stepped to +10 mV for 80 ms (10–90 ms), and returned to baseline for the final 10 ms. Thin black traces represent 500 randomly sampled parameter sets; the thick red trace shows the ensemble mean. Rapid activation is followed by gradual decay, reflecting entry into the inactivated state. Sensitivity analysis was performed on the steady-state window (65–75 ms), during sustained depolarization and active inactivation. Saltelli sampling with base sample size N = 1024 was used for Sobol analysis. (B) First-order (Sᵢ; light bars) and total (S_Ti_; dark bars) Sobol indices averaged over the 65–75 ms window. Error bars denote bootstrap confidence intervals. Parameters are grouped by transition class: C₁–C₂ (closed–closed), C₂–O (activation), and O–I (inactivation). Inactivation transitions dominate steady-state variance, followed by activation transitions, while distal closed–closed transitions remain weak. Prefactor ranges for the O–I transitions were restricted relative to activation transitions to reflect slower inactivation kinetics, yet these transitions account for the largest share of output variance at late times. (C) Interaction contribution for each parameter, quantified as (S_Ti_ − Sᵢ). Interaction effects are concentrated in transitions directly connected to the open state (C₂–O and O–I), whereas distal closed–closed transitions exhibit minimal higher-order contribution. Thus, even when interaction effects are explicitly quantified, the structural suppression of distal transitions persists.

Steady-state Sobol indices reveal a clear redistribution of parameter influence relative to activation-only models. As shown in Figure 5B, the O–I transitions dominate total variance in the late depolarized window (ΣS_T_ᵢ ≈ 0.69), exceeding the contribution of the activation edge C₂–O (ΣS_T_ᵢ≈ 0.44), while the distal closed–closed edge C₁–C₂ remains weak (ΣS_T_ᵢ≈ 0.11). Thus, during sustained depolarization, variance in Pₒ is primarily governed by inactivation kinetics, consistent with the open-state occupancy becoming dynamically limited by exchange with the inactivated state.

Importantly, the inclusion of inactivation does not abolish the structural bottleneck identified in earlier figures. Although the dominant edge shifts from activation to inactivation, parameter influence remains concentrated in transitions directly connected to the open state. The distal C₁–C₂ transitions continue to contribute minimally to steady-state variance.

To determine whether weak first-order indices mask higher-order interaction effects, I examined the difference between total and first-order indices (S_T_ᵢ − Sᵢ, Figure 5C). Interaction contributions are concentrated in the activation and inactivation edges, whereas the distal closed–closed transitions exhibit negligible interaction mass. Thus, the low sensitivity of distal transitions is not rescued by higher-order coupling; it reflects a genuine structural suppression imposed by the linear topology.

Notably, the pre-exponential factor ranges assigned to O–I transitions were narrower than those for activation transitions to reflect slower inactivation kinetics. Despite this conservative sampling range, the inactivation edge dominates steady-state variance. The observed hierarchy, therefore, emerges from the model’s dynamical occupancy structure rather than from permissive prior ranges.

Together, these results demonstrate that adding inactivation redistributes sensitivity toward O–I transitions during sustained depolarization but does not eliminate the structural constraint inherent to linear chains. Even in extended models, parameter influence remains concentrated on edges directly coupled to the open state, while distal transitions remain weak both in first-order and interaction contributions.

To further examine the structural meaning of the sensitivity hierarchy identified in Figure 5, I asked whether the apparent weakness of distal transitions reflects intrinsic irrelevance or simply the relative flexibility of competing bottlenecks. In the baseline C₁–C₂–O–I model (Scheme 4), steady-state variance during sustained depolarization was dominated by the O–I transition, with moderate contribution from the C₂–O transition and minimal contribution from the distal C₁–C₂ transitions (Figure 5B). This hierarchy, however, is evaluated under an ensemble in which all parameters are allowed to vary across their full prescribed ranges.

To test whether the observed sensitivity structure depends on this relative variability, I repeated the ensemble analysis while constraining the C₂–O transition parameters to a narrow range (±1% around their median values), leaving all other parameters sampled across their original ranges. This manipulation effectively reduces the flexibility of the activation bottleneck without altering the model topology.

The resulting redistribution of total Sobol indices is shown in Figure 6A. In the full model (gray bars), parameter influence is concentrated in transitions directly adjacent to the open state, particularly O–I, while the C₁–C₂ transitions exhibit near-zero S_T_ᵢ. In contrast, when variability of the C₂–O transition is constrained (black bars), its contribution collapses and variance is redistributed upstream. The C₁–C₂ transitions, previously weak, become the dominant contributors to output variance.

**Figure 6.**
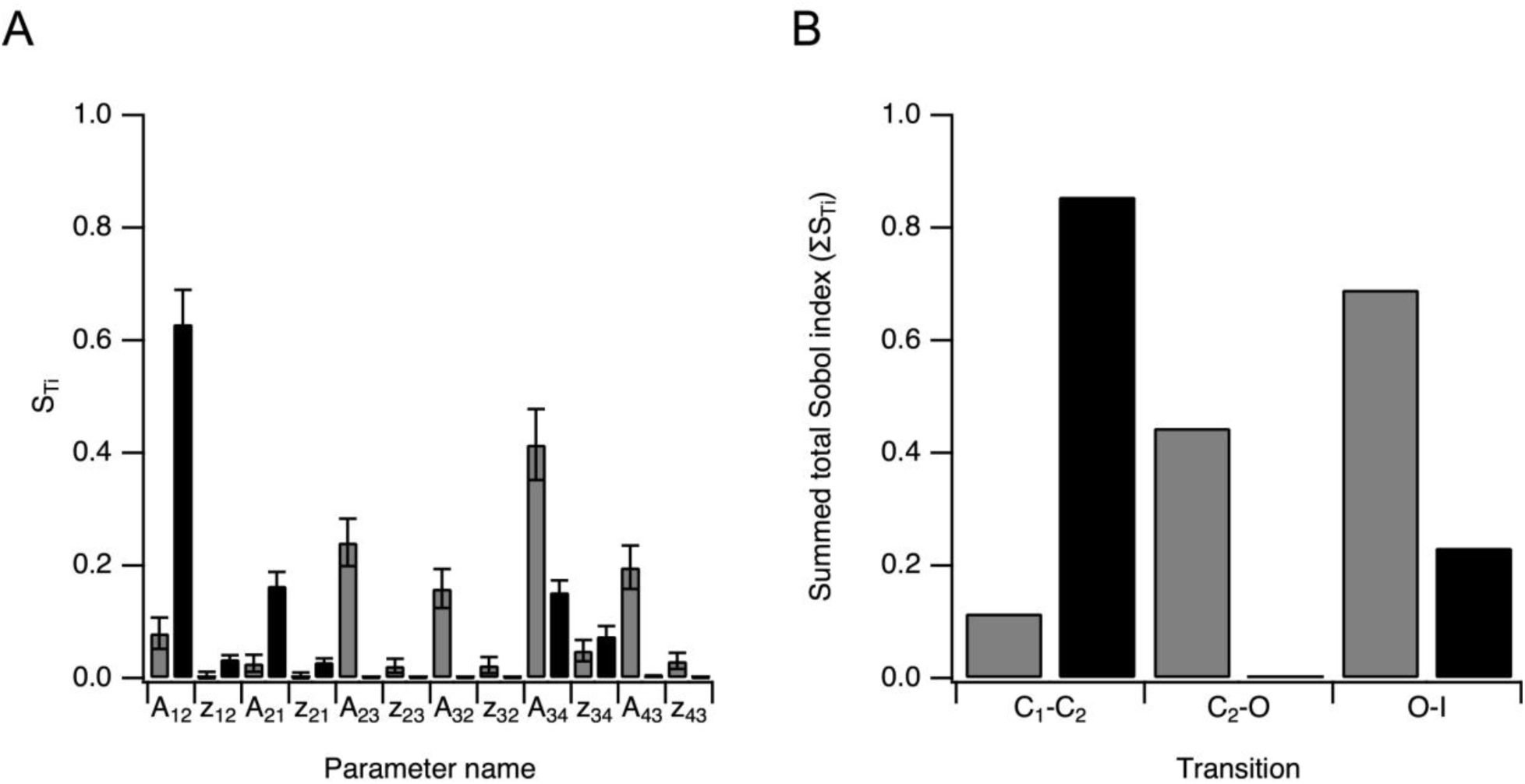
Variance redistribution following constraint of the activation edge. (A) Parameter-level total Sobol indices (STᵢ) for the C₁–C₂–O–I model under sustained depolarization. Gray bars represent the full model in which all parameters were sampled across their original ranges. Black bars represent the constrained model in which variability of the C₂–O transition parameters was restricted to ±1% around their median values, while all other parameters retained their full sampling ranges. In the full model, variance is dominated by transitions adjacent to the open state, particularly O–I, with minimal contribution from distal C₁–C₂ transitions. When variability of the C₂–O edge is constrained, its contribution collapses and variance is redistributed primarily to the upstream C₁–C₂ transitions. (B) Summed total Sobol indices (ΣSTᵢ) aggregated by transition class for the full (gray) and constrained (black) models. In the full model, steady-state variance is dominated by the inactivation edge (O–I), with moderate contribution from C₂–O and minimal contribution from C₁–C₂. Constraining the activation edge shifts the dominant source of variance upstream to C₁–C₂, while the contribution of C₂–O becomes negligible. This redistribution demonstrates that Sobol sensitivity reflects the relative flexibility of competing bottlenecks within the topology rather than the intrinsic structural importance of individual transitions.

Aggregation by transition class (Figure 6B) clarifies this redistribution. In the full model, the inactivation edge (O–I) dominates steady-state variance, followed by C₂–O, with minimal contribution from C₁–C₂. When C₂–O variability is restricted, the dominant contribution shifts to the C₁–C₂ transition, while the C₂–O transition becomes negligible. Thus, variance flows to the most flexible bottleneck within the linear topology.

These results demonstrate that Sobol sensitivity indices quantify the relative contribution of parameters within the context of a specified uncertainty ensemble. Low S_T_ᵢ values do not imply that a transition is structurally unimportant; rather, they indicate that its influence is overshadowed by more variable downstream edges under the chosen sampling scheme. When the dominant bottleneck is constrained, upstream transitions assume control of the output variance. Sensitivity structure, therefore, reflects the interplay between topology and parameter variability, not the intrinsic indispensability of individual transitions.

## Discussion

In this study, I examined the parameter sensitivity of Markov models describing the kinetics of voltage-gated ion channels, aiming to balance model complexity and interpretability. Here, I have shown, using global variance-based Sobol analysis applied to two-state and three-state linear models, a three-state cyclic topology (Scheme 3), and a four-state inactivation model (Scheme 4), that the sensitivity structure is strongly topology-dependent and only weakly protocol-dependent. In linear chains, transitions that are topologically distal from the open state contribute little to the variance of the open probability P_o_ under both step and sinusoidal commands (Figs. 2 & 3). Closing the three-state chain with a direct C_1_-O transition redistributes sensitivity to the newly introduced route and collapses the original bottleneck (Fig. 4), demonstrating that distal-parameter weakness is a consequence of serial topology rather than a universal feature of multi-state gating. Adding an inactivated state shifts dominant variance control to the O-I transition during sustained depolarization while preserving the weakness of distal closed--closed transitions, and explicit decomposition into first-order and total indices shows that this weakness is not rescued by higher-order interactions (Fig. 5). Finally, by constraining variability of the C_2_-O activation transition, I show that variance control can shift upstream to previously negligible transitions (Fig. 6), highlighting that low Sobol indices reflect the relative flexibility of competing bottlenecks within the chosen ensemble rather than intrinsic irrelevance of a transition.

Under standard step voltage protocols, forward transition rates are most influential during activation, whereas reverse transition rates dominate during deactivation (Figs. 1-4). This aligns with the expected biophysical behavior of voltage-gated channels, where occupancy shifts rapidly between closed and open conformations as membrane potential changes. Voltage-dependent parameters (e.g., z_12_, z_21_) show transient sensitivity during activation and deactivation, consistent with their role in shaping the voltage dependence of these shifts. A critical validation condition, stepping to 0 mV, nullifies the exponential voltage term in the rates and drives the corresponding z-indices to zero (Fig. 1D), providing an internal sanity check for the implementation. Such checks are essential in sensitivity workflows that require extensive parameter exploration and are often coupled to stochastic optimization procedures (Keren et al., 2005; Gurkiewicz and Korngreen, 2007; Hay et al., 2011; Ben-Shalom et al., 2012; Ladd et al., 2022).

The quantitative analysis of the three-state linear model makes the structural limitation of serial chains explicit. In this model, the distal C_1_-C_2_ transition remains weak at steady-state activation even when interaction effects are quantified: the interaction contribution S_Ti_−S_i_ is modest and does not elevate weak parameters into dominant contributors (Fig. 2D). This result closes an important methodological loophole: low first-order sensitivity in the linear chain is not simply an artifact of ignoring higher-order coupling. Moreover, extending stimulation from steps to sinusoidal protocols does not change the ordering of parameter importance in any meaningful way. Cycle-averaged S_Ti_ values are stable across 20-200 Hz, and the rank correlation between step-protocol and sine-protocol sensitivity is high (Fig. 3D). This quantitative result establishes directly that oscillatory stimulation does not rescue structurally weak parameters in the serial topology.

The Hodgkin--Huxley formalism can be viewed as a limiting case of simple Markov gating in which all modeled variables are directly tied to transitions that control channel opening, and gating is treated as a concerted mechanism (Hodgkin and Huxley, 1952). Extended Markov chains, in contrast, often introduce long sequences of closed or inactivated states whose transitions are only indirectly connected to opening. The present results show why this is problematic: such transitions can become effectively invisible to macroscopic P_o_ measurements, complicating optimization and inviting parameter inflation without improving predictive power. Importantly, the small interaction contribution in the linear three-state model (Fig. 2D) indicates that invisibility is not simply “hiding” in higher-order effects. At the same time, the bottleneck simulation (Fig. 6) cautions against interpreting low indices as proof of biological irrelevance: when the dominant bottleneck is constrained, control shifts upstream, and previously weak transitions can become the primary source of variance. Thus, in linear chains, distal parameters are weak in the baseline ensemble because variance preferentially flows through the most flexible open-adjacent bottleneck, not because distal transitions can never matter.

The analysis of cyclic topology provides a key mechanistic insight. In the three-state linear chain, all variance must flow through the single open-adjacent C_2_-O transition, creating a bottleneck. Introducing a direct C_1_-O pathway (Scheme 3) creates a parallel route that bypasses the original bottleneck, and the sensitivity hierarchy is fundamentally reorganized: variance shifts to the C_1_-O transition class while the C_2_-O class loses its dominant role (Fig. 4). This mechanistic interpretation is important, because it shows that “distal weakness” is not a generic property of three-state gating; it is a topological consequence of serial arrangement. It also resonates with empirical findings from topology-mutating genetic algorithms, which preferentially generate models in which states are close to the open state and avoid long closed-state chains (Menon et al., 2009). In addition, cyclic topologies naturally impose thermodynamic constraints: microscopic reversibility imposes a constraint per cycle (Colquhoun et al., 2004), reducing the effective parameter dimensionality and improving the conditioning of the estimation problem.

The inactivation analysis (Scheme 4) extends these principles to a physiologically essential feature of voltage-gated sodium channels and many other voltage-gated channels. Under sustained depolarization, sensitivity is dominated by the O-I transition class (Fig. 5B), consistent with open-state occupancy becoming dynamically limited by exchange with the inactivated state on longer timescales. This has practical implications for protocol design and interpretation: sensitivity profiles obtained from short depolarizing steps capture an activation-dominated regime, whereas longer steps spanning the inactivation window reveal a qualitatively different hierarchy. Despite this redistribution toward O-I, the structural bottleneck principle remains intact: distal closed-closed transitions remain weak at late times, and this weakness is not rescued by higher-order interaction mass (Fig. 5C). Thus, adding inactivation does not eliminate the serial-chain limitation; it shifts which open-adjacent transition governs variance in the relevant physiological epoch.

The variance redistribution experiment in Fig. 6 clarifies a subtle but important methodological point. Sobol indices quantify fractional contributions to variance across a prescribed ensemble; they are therefore ensemble-dependent statements about where variability resides, not absolute declarations of indispensability. When the C_2_-O transition is allowed to vary widely, it serves as the primary conduit of variance, and upstream closed--closed transitions contribute little. When variability in the C_2_-O transition is constrained, its contribution collapses and variance shifts upstream to the C_1_-C_2_ transition class (Fig. 6). This result serves as a warning against a naïve reduction rule of the form remove transitions with low S_Ti_. A transition can be weak in one ensemble because it is bypassed by a more flexible downstream bottleneck, yet becomes influential when that bottleneck is fixed by independent measurements, prior constraints, or experimental design. Model simplification is therefore justified only when the redistribution implied by constraining or removing a transition is acceptable for the intended use of the model, not merely because the transition is weak in a particular uncertainty ensemble.

It is also important to distinguish Sobol sensitivity from parameter identifiability. Sobol indices measure how much each parameter contributes to the variance of a specified output across an ensemble; they do not measure whether a parameter can be uniquely inferred from data. A parameter may have low S_i_ yet be identifiable with sufficiently informative data, whereas a parameter with high S_Ti_ may remain practically non-identifiable due to correlations, limited temporal resolution, or insufficiently informative protocols. The distinction between structural and practical identifiability has been clearly articulated in dynamical systems modeling (Raue et al., 2009), and identifiability analysis has been explicitly discussed in ion channel models, and related work has examined fitting strategies and protocol design to improve parameter inference (Fink and Noble, 2009; Clerx et al., 2019). The constructive point of the present work is complementary: for the linear topologies analyzed here, certain parameters contribute so little to the variance of macroscopic P_o_ that they will be poorly constrained by any optimization procedure that relies solely on macroscopic P_o_-based cost functions, because the cost function itself is insensitive to their variation. This amounts to a topology-rooted practical non-identifiability. A natural extension is to incorporate single-channel observables, dwell-time distributions, and transition-frequency statistics, which provide direct access to transitions that are structurally invisible to macroscopic P_o_ (Sakmann and Neher, 1995; Lampert and Korngreen, 2014).

The implications of these findings are twofold. First, they sharpen the limitations of macroscopic recordings and standard protocols in the optimization of complex Markov models. Recent work has shown that information-rich voltage protocols, including sinusoidal voltage commands, can improve the efficiency of ion-channel characterization and parameter fitting (Beattie et al., 2018). In the present analysis, however, extending stimulation from step protocols to multi-frequency sinusoidal commands did not materially alter the sensitivity hierarchy of the linear topology (Fig. 3C & 3D), indicating that richer stimulation alone does not overcome the structural bottlenecks imposed by serial Markov chains. This helps explain why stochastic optimization methods can stagnate in high-dimensional Markov models: low-sensitivity parameters exert minimal influence on the objective function, so convergence is not merely difficult but poorly conditioned (Keren et al., 2005, 2009; Gurkiewicz and Korngreen, 2007; Menon et al., 2009; Hay et al., 2011; Ben-Shalom et al., 2012; Ladd et al., 2022). Second, the results provide practical guidelines for selecting model complexity. Linear chains should be treated with caution: adding distal states increases parameter count without increasing information content in macroscopic P_o_. Cyclic topologies provide a principled alternative by introducing parallel pathways that redistribute sensitivity and by incorporating microscopic reversibility constraints that reduce effective dimensionality (Colquhoun et al., 2004). This conclusion is consistent with topology-search approaches, which tend to avoid long closed-state chains (Menon et al., 2009). Ultimately, bridging the gap between experimental data and computational models remains a central challenge (Nowotny et al., 2007, 2008; Fink and Noble, 2009; Almog and Korngreen, 2014, 2016; Lampert and Korngreen, 2014; Daly et al., 2015; Almog et al., 2022), and the sensitivity maps developed here provide a principled basis for choosing topologies and protocols that maximize the information content of macroscopic measurements. In conclusion, I show that parameter sensitivity in Markov models of voltage-gated ion channels is determined jointly by stimulation protocol, topology, and ensemble definition: oscillatory protocols do not rescue serial bottlenecks, cyclic pathways redistribute control, inactivation shifts dominance to O-I during sustained depolarization without eliminating distal weakness, and apparent insignificance can reverse when a bottleneck is constrained. These results establish practical, topology-sensitive constraints on model complexity and motivate hybrid strategies that combine macroscopic and single-channel data to improve identifiability in biophysically detailed channel models.

## Funding

This work was supported by grants from the Israel Science Foundation to AK (#225/20).

## Author contribution

AK conceived the study, performed the simulations and analyses, and wrote the manuscript.

## Conflict of interest

The author declares no conflict of interest.

## Notes

### Competing Interest Statement

The authors have declared no competing interest.

### Summary of Updates

This revised version substantially expands the scope, rigor, and interpretive clarity of the manuscript. The original submission focused on global Sobol sensitivity analysis in simple linear Markov models of voltage gated ion channels and argued that transitions between closed states not directly connected to the open state contribute little to the variance of the open probability. In the revised manuscript, I retained that central question but extended the analysis in several important directions in order to address reviewer concerns and strengthen the generality of the conclusions. First, the Results section was rebuilt around quantitative steady state summaries rather than relying primarily on time resolved traces. For the three state linear model, I now report first order and total Sobol indices with bootstrap confidence intervals and explicitly quantify the interaction contribution, showing that the weak influence of distal transitions is not rescued by higher order interactions. Second, the sinusoidal stimulation analysis was expanded from a single frequency to a multi frequency comparison at 20, 50, 100, and 200 Hz. This revealed that oscillatory stimulation does not materially alter the sensitivity hierarchy of the linear topology. A direct comparison between step and sine wave rankings showed a strong Spearman correlation, supporting the conclusion that the hierarchy is largely structural rather than protocol specific. Third, I added a full analysis of a cyclic three state topology in which a direct closed to open transition closes the original linear chain into a loop. This demonstrates that the apparent weakness of distal parameters in the linear model is not a universal property of three state Markov schemes, but rather a consequence of serial arrangement. Once the loop is closed, sensitivity is redistributed to the new shortcut pathway. Fourth, I introduced a four state linear model including inactivation. Under sustained depolarization, the dominant source of variance shifts from activation to the open to inactivated transition, yet distal closed to closed transitions remain weak, and this weakness is again not rescued by interaction effects. Finally, I added a variance redistribution experiment in which variability of the dominant activation edge was strongly constrained while all other parameters remained free. Under these conditions, previously negligible upstream transitions became the dominant contributors to output variance. This result clarifies that low Sobol indices reflect the relative flexibility of competing bottlenecks within the sampled ensemble rather than the intrinsic irrelevance of a transition.

